# Exceptional longevity of mammalian ovarian and oocyte macromolecules throughout the reproductive lifespan

**DOI:** 10.1101/2023.10.18.562852

**Authors:** Ewa K. Bomba-Warczak, Karen M. Velez, Luhan T Zhou, Christelle Guillermier, Seby Edassery, Matthew L. Steinhauser, Jeffrey N. Savas, Francesca E. Duncan

**Affiliations:** Department of Neurology, Northwestern University Feinberg School of Medicine, Chicago, IL; Department of Obstetrics and Gynecology, Feinberg School of Medicine, Northwestern University, Chicago, IL; Department of Medicine, Aging Institute, University of Pittsburgh School of Medicine, Pittsburgh, PA; Department of Medicine, Division of Genetics, Brigham and Women’s Hospital, Boston, MA

## Abstract

The mechanisms contributing to age-related deterioration of the female reproductive system are complex, however aberrant protein homeostasis is a major contributor. We elucidated exceptionally stable proteins, structures, and macromolecules that persist in mammalian ovaries and gametes across the reproductive lifespan. Ovaries exhibit localized structural and cell-type specific enrichment of stable macromolecules in both the follicular and extrafollicular environments. Moreover, ovaries and oocytes both harbor a panel of exceptionally long-lived proteins, including cytoskeletal, mitochondrial, and oocyte-derived proteins. The exceptional persistence of these long-lived molecules suggest a critical role in lifelong maintenance and age-dependent deterioration of reproductive tissues.

**One sentence summary:** Exceptionally long-lived macromolecules in mammalian ovaries and oocytes as pillars for lifelong reproductive health span.

## Main Text

The female reproductive system is the first to age in the human body with fertility decreasing for women in their mid-thirties and reproductive function ceasing completely at menopause (*1*). In the ovary, aging is associated with a loss in gamete quantity and quality which contributes to infertility, miscarriages, and birth defects (*2-5*). Moreover, the age-dependent loss of the ovarian hormone, estrogen, has adverse general health outcomes (*6*). These sequelae are significant as women globally are delaying childbearing and the gap between menopause and lifespan is widening due to medical interventions (*7, 8*). Although aging is a multifaceted process, loss of proteostasis and dysfunctional protein quality control pathways are hallmarks of reproductive aging (*9*).

Proteostasis relies on tight inter-regulation of protein synthesis, post-translational modifications, folding, and degradation (*10*). While most protein lifetimes in mammals fall within the scale from hours to days (*11, 12*), a subset of intracellular proteins persists for months in rodents (*13, 14*). These long-lived proteins (LLPs) are enriched in tissues harboring long-lived post-mitotic terminally differentiated cells, such as the brain and heart (*13-15*). Although the extended lifespan of LLPs places them at inherent risk for accumulating damage during aging, many of them provide key structural support for the lifelong maintenance of highly stable protein complexes in cells (*16*).

The mammalian ovary is comprised of a fixed and nonrenewable pool of long-lived cells or oocytes. In humans, oocytes initiate meiosis during fetal development, and by birth, all oocytes are arrested at prophase of meiosis I (*17, 18*). This cell cycle arrest is maintained until ovulation, which occurs any time between puberty and menopause, and thus can span decades. The oocytes are particularly sensitive to protein metabolism alterations because they contribute the bulk cytoplasm to the embryo following fertilization. Thus, maternal proteins produced during oogenesis are essential to generate high-quality gametes (*9*). The ovarian microenvironment is a critical determinant of gamete quality and has been shown to become fibro-inflamed and stiff with age (*19-21*). Although a small number of oocyte-specific proteins have been identified as long-lived, including cohesins and several centromere-specific histones, there has not been a discovery-based approach to define the long-lived proteome of the ovary and oocyte. Thus, the potential contribution of LLPs to the age-related deterioration of the reproductive system in mammals remains to be elucidated. In this study we used multi-generational whole animal metabolic stable isotope labeling and leading mass spectrometry (MS)-based quantitative proteomic approaches to visualize and identify ovarian and oocyte long lived macromolecules in *vivo* during milestones relevant to the reproductive system.

### Exceptional longevity of ovarian structures and molecules in mammals

The mammalian ovary is a structurally complex, heterogenous, and dynamic organ with follicles at different stages of development, remnants of ovulation (corpora lutea), and a heterogeneous stroma (*22*) (**Fig. 1A**). Very little is known about the long-term homeostasis and relative turnover of the ovarian tissue components during aging. To address this, we visualized the lifespan of ovarian macromolecules in mammals using a combination of stable isotope labelling and multi-isotope imaging mass spectrometry (MIMS) **(fig. S1A).** First, using a two-generational metabolic labelling of animals with ^15^N, we generated a cohort of fully ^15^N-labelled pups. After birth, labelled females were kept with the labelled dam until weaning, at which time their food source was switched to the ^14^N-chow chase. Using previously established methods (*9, 23-25*), we determined that chase periods of 6 and 10 months represented a biologically relevant reproductive aging continuum as mice within this age range begin to manifest ovarian aging phenotypes, including follicle loss (**fig. S1 B-D)**, decreased ovulation (**fig. S1 E and F)**, and increased fibrotic foci in the ovarian stroma (**fig. S1 G-J**). Importantly, sufficient numbers of oocytes can still be collected at these timepoints for meaningful further downstream analyses of this rare cell type. We first performed MIMS on ovarian sections to visualize and quantify the abundance of ^14^N, representing molecules which have been replaced during the chase period (blue-teal), and ^15^N, which represents ^15^N-containing molecules that must have persisted through the chase period and therefore are long-lived (orange-pink) (**Figs. 1B, S2 A and B**) (*26-28*). Within the ovarian follicles, MIMS revealed a strikingly higher abundance of ^15^N containing molecules in primordial and primary stages relative to later stage follicles, suggesting that primordial follicles can persist for months with limited macromolecular turnover (**Fig. 1C)**. As follicles progress through the primary, secondary, and antral stages the ^15^N/^14^N ratio decreases due to signal dilution associated with the increase of cell number and follicle growth (**Figs. 1C, S2 C**). This change in ^15^N abundance was most apparent in the granulosa cells where long-lived molecules were significantly higher in those within primordial and primary follicles compared to later follicle stages (**Fig. 1C** and **F**). In addition to granulosa cells, long-lived molecules also localized to the basement membrane of some early growing follicles (**fig. S1D**). Beyond the follicle, other somatic compartments of the ovary were additionally found to have higher ^15^N:^14^N ratio suggesting enrichment in long-lived components, including steroidogenic cells (theca layer and corpora lutea), stromal cells, and cells within the ovarian surface epithelium (OSE) (**Fig. 1 B** and **D**). Our quantitative analysis revealed that the OSE had significantly higher ^15^N/^14^N ratio among the mentioned cell types (**Fig. 1G**). Lastly, we analyzed the ^15^N/^14^N ratios in the nucleus relative to the cytoplasm, which revealed significant enrichment of long-lived, ^15^N-positive molecules in the nuclei of granulosa cells within primary follicles, and cells within the corpora lutea, the stroma, and the OSE. (**Fig. 1 E** and **H**). As both proteins and nucleic acids contain nitrogen, the ^15^N-positive nuclear signal could correspond to known long-lived nuclear proteins, such as histones, nuclear pore proteins, and lamins, which were previously identified in neuronal, post-mitotic cells (*14, 15*). Alternatively, as these nuclear ^15^N-hotspots coincided with the ^31^P signal, which is enriched in DNA and correlates with DNA labeling(*29*), this data may suggest that the DNA itself is long-lived (**Fig. 1E**). These results reveal that distinct macromolecular components within select ovarian cells and tissue regions persist throughout the healthy reproductive stage, with limited renewal, and those long-lived molecules persist through the stage where ovaries manifest marked reproductive aging phenotypes.

**Fig. 1:**
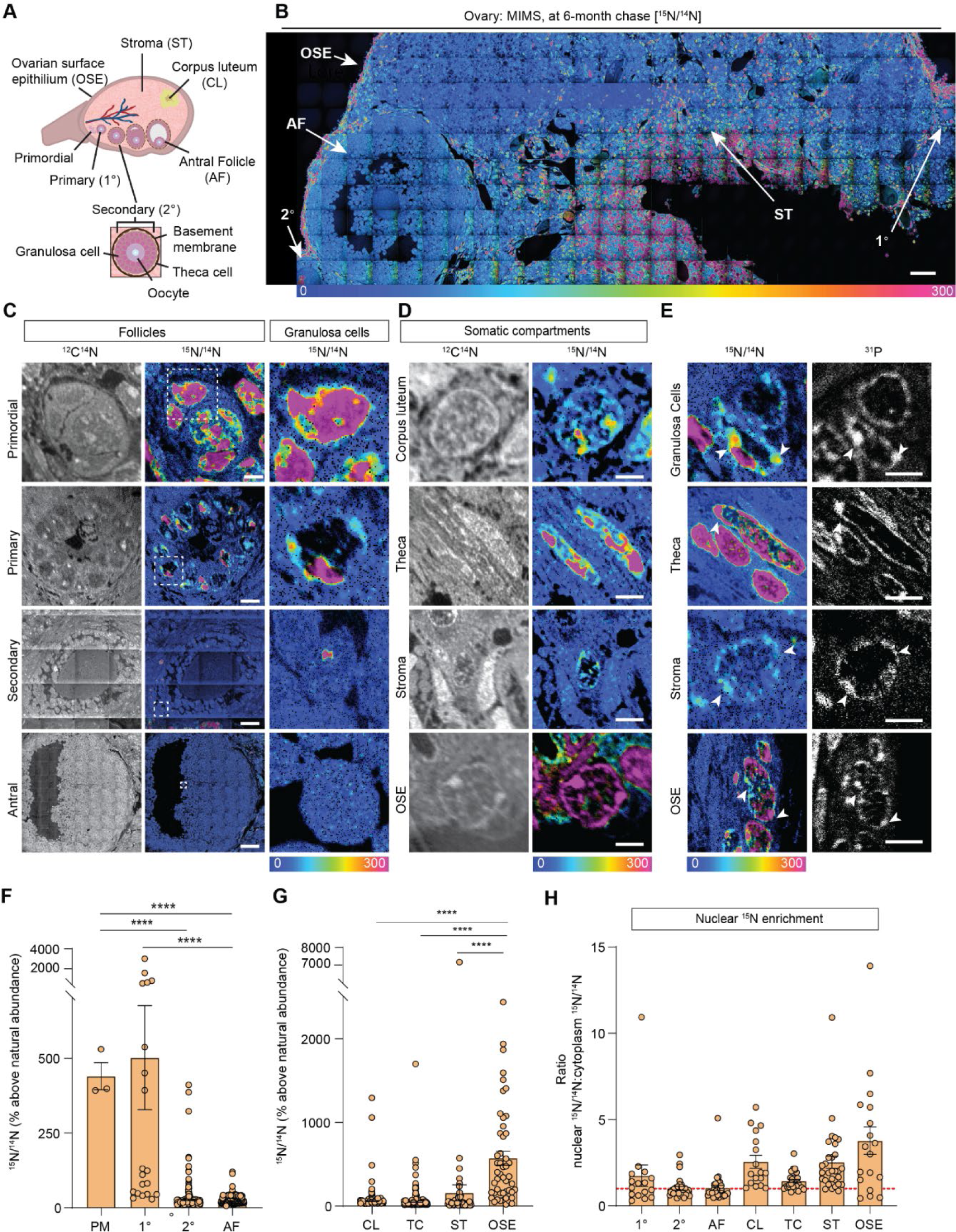
MIMS analysis of cells and structures across ovarian tissue sections. **(A)** Diagram depicting and defining structures of the mammalian ovary. **(B)** Representative hue saturation intensity (HSI) mosaic from an ovary of a ^15^N labeled mouse (6 months old). Localization and abundance of ^15^N varied depending on cell type. HSI scale was set to 0% (natural ^15^N/^14^N ratio) to 300% (above the natural ratio). Scale bar = 60 µm. **(C)** High abundance of ^15^N is seen in early-stage follicles, specifically within granulosa cells. **(D)** Representative images of somatic cells show differences in ^15^N labeling. **(E)** Intracellular abundance of ^15^N is colocalized with ^31^P abundance across all cell types. **(F)** Differences in ^15^N/^14^N ratios reveal granulosa cells of early-stage follicles have greater ^15^N abundance than later stages. **(G)** Among somatic cells, quantitative analysis shows a greater abundance of ^15^N at the ovarian surface epithelium. **(H)** Ratio analysis show abundance of ^15^N localized in nuclear regions of cells. A hypothetical ratio of one, denoted as a red dash line, signifies no difference in ^15^N abundance between cytoplasmic and nuclear regions. Abbreviations: PM (primordial follicle), 1° (primary follicle), 2° (secondary follicle), AF (antral follicle), CL (corpus luteum), TC (theca cell), ST (stroma), and OSE (ovarian surface epithelium). HSI scale for all images was set to 0%-300% (above natural abundance). Data are shown as mean±SEM. Statistical analysis was performed using a one-way ANOVA. Asterisk denotes statistical significance (* p≤.05; ** p≤.01; *** p≤.001; **** p≤.0001). Scale bar = (B): 50 µm; (C): 3 µm (PR), 6 µm (1°), 30 µm (2°), 75 µm (AF); (D): 2.5 µm (CL), 2.5 µm (TC), 5 µm (ST), 2.5 µm (OSE); (E): 3 µm (GC), 3 µm (TC), 2.5 µm (ST), 6 µm (OSE).

### Identification of the long-lived proteome in mammalian ovaries

Although MIMS analysis provides important spatial information on long-lived structures in the ovary, it does not provide the identity of ^15^N-containing macromolecules that comprise them. To address this, we performed liquid chromatography mass spectrometry (LC-MS/MS)-based proteomic analysis of ovarian tissues isolated from metabolically labeled mice. After 6-months of ^14^N-chase, we identified 36,222 ± 6768 ^14^N-peptides mapping to 4106 proteins, and 13 ± 5 ^15^N peptides, which collectively mapped to 33 LLPs across all the biological replicates (**Fig. 2 A and B, table S1**). To gain a deeper insight into the persistence of LLPs beyond 6 months, we also analyzed ovaries isolated from females that remained on ^14^N chase for 10 months. Although the total numbers of both ^14^N and ^15^N peptides and proteins were similar between the two timepoints (39,637 ± 12005 ^14^N-peptides mapping to 4464 proteins and 13 ± 6 ^15^N peptides mapping to 15 LLPs), the majority of LLPs identified at 6 months were no longer identified as long-lived at the 10-month chase timepoint. Only tubulins and select histones persisted and were identified as LLPs at this aged timepoint. Gene ontology (GO) enrichment analysis of LLPs identified at the 6-month chase timepoint revealed significant overrepresentation for terms related to chromatin, nucleosome, tubulin complex, and mitochondria (**Fig. 2C**).

**Fig. 2.**
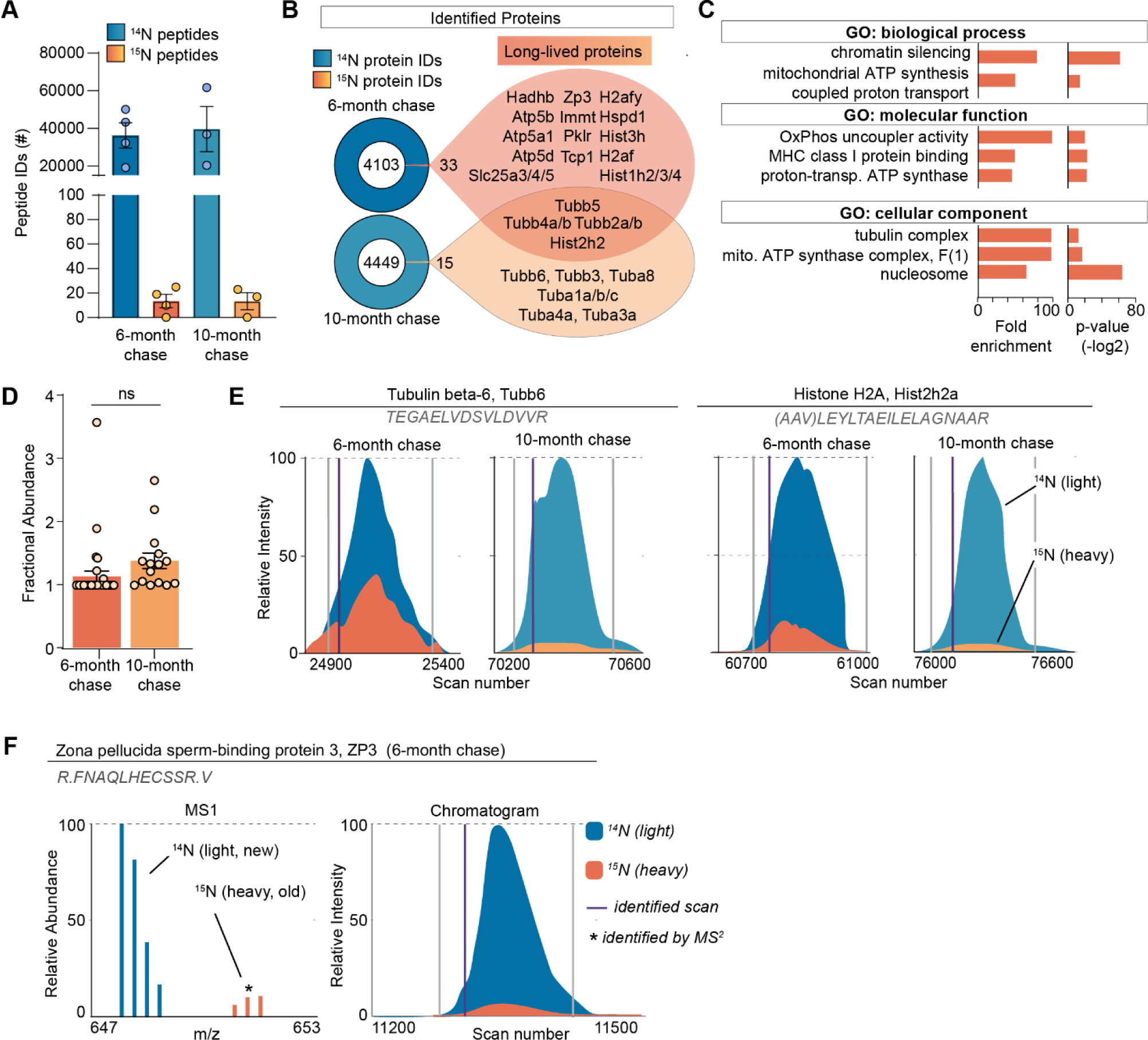
Long-lived proteome in mammalian ovaries. **(A)** Summary of peptide identification at 6 and 10-month chase points, blue graphs indicate ^14^N peptide IDs, orange/yellow graphs indicate ^15^N peptide IDs. **(B)** Summary of protein identification at each time point, with a list of proteins identified as long-lived in orange. **(C)** GO analysis of the LLPs identified in ovaries at 6-months chase revealed that terms related to chromatin, nucleosome, tubulins, and mitochondria are significantly enriched. **(D)** Fractional abundance of LLPs identified at 6 and 10-month chase. **(E)** Annotated representative chromatograms of two representative proteins that persist through both 6- and 10-montch chase, illustrating the decreasing ^15^N-signal over time. blue - 14N (new), orange 15N (old), purple line: identified scan. (**F)** Annotated representative raw MS1 scan of Zona Pellucida protein 3 (ZP3). Mean ± SEM; 3-4 biological replicate per timepoint, ns – not significant by Student’s t-test.

Next, we determined the quantity of each LLP remaining after the ^14^N-chase period by calculating the fractional abundance (FA; ^15^N remaining, ^15^N/[^14^N + ^15^N]) for each LLP in the ovary using reconstructed MS1 chromatograms from LC-MS/MS analysis (*30*) (**Fig. 2 D-F**). We found that LLPs had significant differences in FA between the 6- and 10-month time points, with 1.13 ± 0.08 % and 1.37 ± 0.12 % ^15^N-remaining, respectively (**Fig. 2D**). The discrete pool of proteins that persisted throughout both time points allowed for a unique direct comparison of ^15^N to ^14^N-peptide peak intensities. This analysis showed reduced abundance of ^15^N at the 10-month chase timepoint, consistent with continual, albeit slow, turnover of the protein pool (**Fig. 2E**). Interestingly, at the 6-month chase timepoint, we identified ^15^N peptides mapping to an oocyte-specific protein, Zona pellucida-3 protein (ZP3), indicating that a pool persists without turnover for at least 6 months, but less than 10 months, as long-lived ZP3 was no longer identified at the 10-month timepoint (**Fig. 2F**). While histones and tubulin have been previously identified as LLPs in the mammalian brain and heart tissues, the 6-month long persistence of ZP3 in mouse ovaries is unexpected and of potential importance to reproductive biology.

### Exceptional longevity of mitochondrial and myosin proteins in mammalian oocytes

MIMS analysis of ovarian sections also captured oocytes at various stages of development, which in addition to the enrichment of ^15^N-signal within the oocyte nucleus, revealed multiple smaller cytoplasmic ^15^N-hot spots (**Fig. 3A**). Thus, to determine the identity of these ^15^N-enriched molecules we isolated fully grown oocytes from ovaries of labeled mice at 6 and 10 months chase timepoints followed by LC-MS/MS analysis (**Fig. 3B and S3A**). In oocytes, we identified a total of 2919 proteins at 6-months and 3234 proteins at 10-months (**Fig. 3B, table S2**). Although after 6-months of chase, we identified 146 LLPs in oocytes, only 11 LLPs were identified after 10-months, indicating that by this timepoint a vast majority of LLPs have been degraded and renewed. Interestingly, the GO analysis of LLPs in oocytes identified at 6-month timepoint revealed a significant enrichment of terms related to nucleosomes, myosin complex and several additional terms related to mitochondria including - OxPhos complexes, mitochondrial nucleoid, TCA cycle complexes, and mitochondrial permeability transition pore complex (**Fig. 3B**).

**Fig 3.**
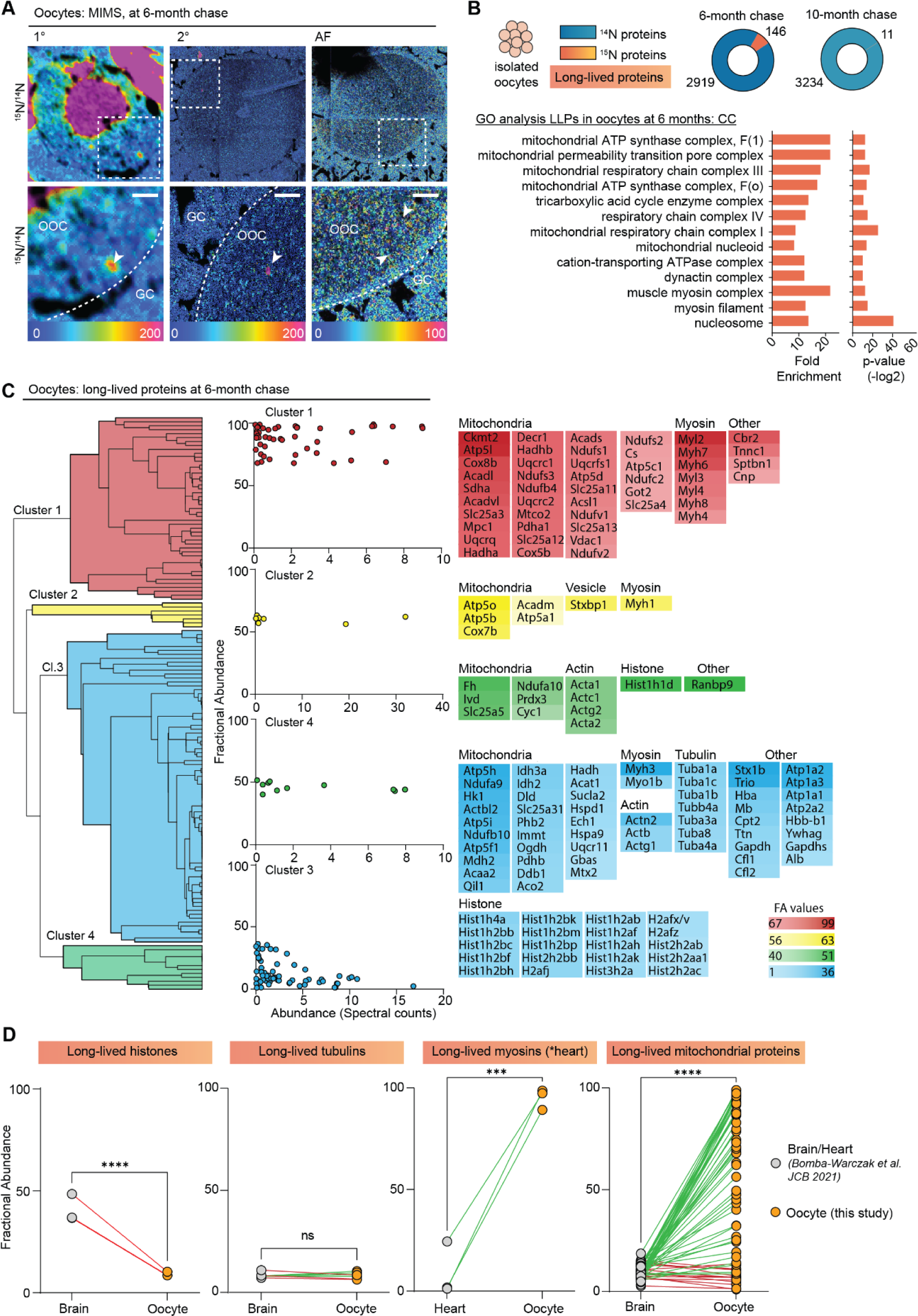
Exceptional longevity of nuclear, cytoskeletal, and mitochondrial proteins in mouse oocytes. **(A**) MIMS analysis reveals high abundance of ^15^N in nuclei and throughout the cytoplasm of oocytes. **(B)** Purified oocyte population was harvested from pulse-chased mice and analyzed using gel-LC-MS/MS. Charts illustrate the number of proteins identified at each timepoint (blue) along with long-lived proteins (orange). GO analysis of the LLPs identified in oocytes at 6-months chase revealed an enrichment for terms related to nucleosome, myosins, and mitochondria. **(C)** Hierarchical cluster analysis of fractional abundance of LLPs identified in oocytes. **(D)** Direct comparison of fractional abundances of proteins previously identified as long-lived and LLPs in oocytes. Mean ± SEM; Oocytes collected from 4-7 females per time-point, *** p-value <.001, **** p-value < .0001 by Kruskal-Wallis ANOVA with Tukey’s multiple comparisons test. Scale bar = 4 µm.

Next, we quantified the fraction of each protein pool that persisted for 6 or 10 months by calculating fractional abundance (FA) values, where the higher the value the longer-lived the corresponding proteins are (**fig. S3B**). Overall, there was no significant difference in FA between the two chase timepoints, with 41 ± 2.9 and 50.8 ± 12.2 ^15^N-remaining at 6 and 10-months, respectively (**fig. S3C**). The higher average FA average at 10-month chase is likely due to turnover of proteins with lower FA values at 6 months, which by 10-months would leave the oocyte with the most persistent pool of proteins. In agreement, the FA values for 4 LLPs that were identified at both time points (Hba, Atp5a, Atp5B, and Hist1h4a) sharply decline between 6 and 10 months (**fig. S3D**), demonstrating protein degradation and replenishment. Considering the high abundance of mitochondrial proteins in our data set, and recent reports showing that expression of mitochondrial proteins is suppressed in aging oocytes (*31*), we compared the number of mitochondrial proteins and spectral counts identified (**fig. S3E and F**), as well as fractional abundance (**fig. S3G**) for mitochondrial proteins across the two timepoints. Our results indicate that while there are no significant differences in total mitochondrial protein abundance at 6- and 10-months post-chase, the pool of long lived mitochondrial proteins decreased significantly, with the majority of proteins being turned over at the later timepoint.

Hierarchical clustering of the LLPs identified at the 6-month timepoint revealed mitochondrial proteins and myosins as the two protein groups with the highest fractional abundance in the oocyte (**Fig. 3C**). In particular, mitochondria exhibited a wide range of FA values ranging from 1.10% to 98.9%, with an average of 55.9 ± 35.1 of ^15^N-remaing at 6-months (**table S2**). FA values for myosins was markedly higher than both actin and tubulin with an average FA value of 80.97 ± 19.8 % for myosins, 31.4 ± 15.6% for actins, and only 8.3 ± 1.4% for tubulins. This indicates that while all three cytoskeletal components are long-lived, nearly 81% of the myosin protein pool persists throughout the 6-month timepoint, whereas only 31% of actin protein pool and 8.3% of tubulin protein pool persists throughout the same length of time. Histones were also identified as LLPs in oocytes with average FA values of 7.7 ± 9.8. Mitochondrial proteins, histones, myosins, and tubulins have been previously identified as LLPs in brain and heart tissues, which are known to contain long-lived terminally differentiated cells (i.e. neurons in the brain and cardiac myocytes in the heart) (*13*). Importantly, however, this is the first time that a subset of the same proteins has been identified as long lived in the germ cell (**Fig. 3D**). Interestingly, the fractional abundance of mitochondrial LLPs in the brain (10.2 ± 6.6%)(*13*) and myosins in the heart (4.6 ± 8.1%)(*13*) is much lower in than the same LLPs quantified in the oocyte (55.9 ± 35.1)(**Fig. 3D**). In contrast, histones were less long-lived in the oocyte compared to the brain, and there was no significant difference observed for tubulins (**Fig. 3D**). Thus, although the identity of long-lived proteins may be conserved across tissues with long-lived cells, differences in fractional abundance may reflect tissue specific functions and requirements.

## Discussion

In this study, multi-generational whole animal metabolic stable isotope labeling, paired with multi modal MS-based quantitative approaches enabled visualization and identification of ovarian and oocyte long lived macromolecules in vivo. Our findings provide a novel framework for how long-lived structures may regulate gamete quality. Long-lived macromolecules localized throughout the ovary including the follicular compartment with prominent signals in the granulosa cells of primordial and primary follicles relative to later stage growing follicles. These findings are consistent with the knowledge that the squamous pre-granulosa cells surrounding the oocyte within primordial follicles form early in development which coincided with the ^15^N-labelling pulse period. These squamous granulosa cells are generally thought to lack the ability to undergo mitotic division until follicles are activated to grow, so it is not surprising that we observed long lived macromolecules persisting within them (*32*). In contrast, granulosa cells in growing follicles are generated by cell divisions that take place during follicle activation and growth which coincided with the ^14^N-chase period. Thus, long-lived structures (i.e. enriched with ^15^N) were diluted through cell divisions during follicle growth. Moreover, during follicle growth, granulosa cells proliferate and differentiate adding new pools of synthesized proteins and molecules. Our results demonstrate that macromolecules formed early in development can persist in squamous granulosa cells for months. Thus, it is possible that these long-lived molecules will accumulate more damage in primordial follicles that remain quiescent for longer periods relative to those that activate earlier. Whether such damage occurs and how it translates into decreased follicle survival or gamete quality will require further investigation.

Within the extrafollicular ovarian environment, the OSE exhibited a striking enrichment of long-lived molecules. The OSE is highly dynamic due to repeated post-ovulation wound healing and repair, and its regenerative capacity occurs through a somatic stem/progenitor cell-mediated process (*33*). Interestingly, long-lived proteins are retained in other cells undergoing repeated asymmetric divisions and are speculated to contribute to the reproductive aging process (*34*). Consistent with this possibility, the architecture and wound healing ability of the OSE is altered with advanced reproductive age (*35*). Furthermore, nuclear enrichment of the ^15^N signal was highest in cells of the OSE. It is plausible that the older template DNA is segregated into the daughter cell destined to become the stem cell to ensure genetic stability of the OSE. Better understanding of the dynamics of long lived molecules in the OSE will require generation of specific samples at precise stages of the estrous cycle and across a time course of ovulation to capture follicular rupture and repair.

Through LC-MS/MS analysis, we identified specific long-lived proteins in the mammalian ovary across the reproductive lifespan. LLPs tend to be part of large protein complexes and include histones, nuclear pore complex proteins, lamins, myelin proteins, and mitochondrial proteins (*16*). In the ovary, the major categories of LLPs included histones, cytoskeletal proteins, and mitochondrial proteins. ZP3 was an oocyte-derived protein identified to be long-lived for at least 6 months. ZP3 is a protein that comprises the zona pellucida (ZP) or glycoprotein matrix of the oocyte, and it is expressed in oocytes of actively growing follicles beginning at the primary stage when the ZP begins to form (*36*). However, during the pulse period, there would have been very few growing follicles in the ovary because of the immature age of the mice, and most importantly, none of these follicles would have persisted 6 months since folliculogenesis only takes approximately 21 days (*37*). These findings suggest that ZP3 may be expressed earlier in oocyte development than previously anticipated. Because LLPs can be at the core of scaffold complexes, a primitive zona may exist at the primordial follicle stage upon which the bona fide ZP is established in growing follicles (*38, 39*). Consistent with this, expression of ZP proteins has been observed in human primordial follicles (*39*). Interestingly, there are documented age-related defects in the structure and function of the ZP which occur with time-dependent scaffold deterioration. An alternate explanation for our observation is that ZPs from atretic follicles persist and become incorporated into the ovarian matrix. Precedent for this exists because ZP proteins have been identified as components of the matrisome of decellularized porcine ovaries (*40*). Interestingly we did not identify ZP3 as an LLP in isolated fully grown oocytes which provides further support that in the ovary, the long-lived pool of ZP3 is derived from primordial follicles or are within the matrix. These possibilities require further investigation and may not be mutually exclusive.

Most LLPs were degraded and replaced between 6-10 months of chase. At the 6 month time point, we detected more long lived proteins than the 10 month time point in both the ovary and the oocyte because proteins degrade over time, and more time has elapsed at the later time point. Moreover, between the 6 and 10 month time points, age-related tissue dysfunction is already evident in the ovary. For example, in 6-9 month old mice, there is already a deterioration of chromosome cohesion in the egg which results in increased interkinetochore distances (*41*), and by 10 months, there are multinucleated giant cells present in the ovarian stroma which is consistent with chronic inflammation (*20*). Thus, our results suggest that important shifts in the proteome occur during mid to advanced reproductive and may be another early feature of ovarian aging. Whether the LLPs at the 6 month time point serve as a protective mechanism in maintaining gamete quality or whether they contribute to decreased quality associated with reproductive aging is an intriguing dichotomy which will require further investigation.

A small subset of tubulins and histones persisted throughout the entire 10-month chase period, indicating that their replacement is exceptionally slow. LLPs which were present at both time points and persisted to at least 10 months may have important roles in the aging process. Interestingly, Tubb5 and Tubb4a have high homology to primate-specific Tubb8, and Tubb8 mutations in women are associated with meiosis I arrest in oocytes and infertility (*42, 43*). Thus, perturbation of these particular proteins by virtue of their long-lived nature may be associated with impaired function and poor reproductive outcomes, and these possibilities warrant future investigation. We noted that LLPs identified at 10 months were not always identified as long-lived at 6 months. This is a common limitation of mass spectrometry-based proteomics where each sample is prepared and run individually, which introduces variability between biological replicates, especially with respect to low abundant proteins. To compensate for this known and inherent variability, we applied stringent filtering criteria where we required long-lived peptides to be identified in an independent MS scan which provided us peptides of highest confidence.

By isolating fully grown oocytes from the ovary, we were able to determine the long-lived proteome of a purified germ cell population across the reproductive lifespan. Although we identified certain histone proteins as long lived, their relative fractional abundance was much lower than in the brain, a tissue which also contains post-mitotic cells (*14, 15*). However, histone-variant exchange occurs continuously during mouse oogenesis and is required for both transcriptional regulation and de novo DNA methylation (*44*). Thus, turnover and exchange of histones is likely more dynamic than previously assumed in terminally differentiated or post-mitotic cells, and our findings are consistent with this in the oocyte. Myosin and actin were also identified as LLPs with relatively high fractional abundance indicating exceptional longevity beyond 6-months. These proteins play numerous roles in oocyte maturation, fertilization, and egg activation, including nuclear positioning, spindle rotation and anchoring, chromosome segregation, cytokinesis, cortical granule exocytosis, and cytoplasmic flow (*19, 45-49*). Defects in many of these processes have been reported with advanced reproductive age, but whether long-lived pools of actin and myosins contribute to this etiology remains to be elucidated (*50*). Interestingly, F-actin stabilization restricts chromosome segregation errors due to cohesion loss which increase with age, so long lived pools of actin may confer a beneficial effect and protect against aneuploidy (*49*). The molecular mechanism(s) governing the longevity of the specific identified proteins in oocytes and ovaries as well as their relationship to the age-related decline in fertility and ovarian function require further investigation.

Mitochondrial proteins were the predominant LLPs in isolated oocytes across the reproductive lifespan, and it is plausible that ^15^N cytoplasmic hotspots in oocytes that were observed in MIMS may correspond to mitochondria. Various aspects of mitochondrial dysfunction have been long-implicated in the age-dependent decline in gamete quality, with an age-dependent decrease in the number of mitochondria, an increase in abnormal morphology, and altered subcellular distribution (*51*). Moreover, mtDNA copy number decreases with age, whereas mtDNA mutations increase (*52, 53*). Finally, the functional capacity of mitochondria decreases with age with decreased membrane potential, increased reactive oxygen species, increased oxidative stress, and decrease energy output (*54*). Although it is possible that oocyte long lived mitochondrial proteins deteriorate with age and contribute to mitochondrial dysfunction, a different model is emerging for these proteins. Mitochondrial proteins are exceptionally long lived in tissues containing long-lived terminally differentiated cells and have now been documented in the brain (neurons), heart (cardiac myocytes), and the ovary (oocytes) (*13*). These mitochondrial LLPs are primarily localized to the cristae invaginations, which are throughout to serve as long-term stable ultrastructure within mitochondria. This strategic enrichment is hypothesized to serve as a lifelong structural pillar of mitochondria to support and maintain these organelles over long time frames (*16*). Although mitochondrial LLPs persist for at least 6 months in oocyte, the majority were undetectable by 10 months. Thus, it is tempting to speculate that a stable pool of mitochondrial LLPs provides structural support for the maintenance of mitochondrial structure and function early during the reproductive lifespan. However, turnover of mitochondrial LLPs later in the reproductive lifespan may serve as a biological timer of aging. Because mitochondria are maternally inherited, these mitochondrial LLPs formed during fetal development of the mother are likely transferred to the embryo and impact subsequent generations (*55*). Further investigation into the role of long-lived mitochondrial proteome in oocytes is necessary to understand their life-long contribution to reproductive health and fertility outcomes.

## Author contributions

Conceptualization: EKBW, JNS, FED; Methodology: JSN, FED, EKBW, KMV, LTZ, SE, CG, FG, MS, Investigation: EKBW, KMV, LTZ, SE; Visualization: EKBW, KMV; Funding acquisition: JNS, FED; Project administration: JNS, FED; Supervision: JNS, FED; Writing – original draft: EKBW, KMV, JNS, FED; Writing – review & editing: all authors.

## Data and materials availability

All data generated or analyzed during this study are included in the manuscript and supporting files. RAW MS data has also been deposited at MassIVE under the accession number MSV000092217

## Supplementary Materials

### Materials and Methods

#### Animals

Mice of the FVB strain were obtained from Jackson Laboratory (Bar Harbor, ME). For the whole animal isotope pulse-chase labeling strategy, female (5 weeks old) and male mice (10 weeks old) mice were obtained. To validate the biological relevance of our chase period, reproductive aging parameters were evaluated in unlabeled female mice at 6 weeks, 6 months, and 10 months of age. Retired female FVB mouse breeders were obtained at 15 weeks of age and aged out to 6 months and 10 months. Mice were acclimated in the vivarium upon arrival for at least two weeks prior to experimental use. All mice were housed at Northwestern University’s Center for Comparative Medicine under constant temperature, humidity, and light (14 hours light/10 hours dark). Mice were fed and provided with water *ad libitum*. All animal care and experimental protocols in this study were conducted under the guidelines set by the NIH Guide for the Care and Use of Laboratory Animals handbook, and the animal protocol was approved by the Animal Care and Use Committee of Northwestern University.

#### Pulse-chase labeling strategy

Mice were metabolically labeled using a two-generation metabolic pulse-chase labeling strategy as previously described (*13*). Briefly, three female FVB mice were fed a spirulina ^15^N-containg chow for the duration of the study (initial labeling, breeding, pregnancy, and weaning) (Cambridge Isotope Laboratories, Inc., Tewksbury, MA). After 13 weeks of being on a ^15^N diet, these labeled females were co-housed with unlabeled males for 5 days to allow a complete estrous cycle for breeding. Pregnant females were housed separately to allow for accurate dating of litter births. All females were allowed to breed a total of 3-5 times to obtain sufficient pups for the chase period. The pulse period was defined as the timespan between gestation and weaning of pups. Thus, pups conceived from labeled females had nitrogen-containing molecules and proteins labeled with ^15^N. At 22 days old, pups were weaned and fed a ^14^N diet *ad libitum*. Pups were maintained on a ^14^N diet (chase) until they reached 7 and 11 months of age at which point tissues were harvested for downstream analyses. This animal labeling strategy generated 19 female pups. For ovary studies, 4 mice were used at 6 months post-chase and 4 mice at 10 months post-chase. One ovary per mouse was used for MIMS, and the contralateral ovary was used for LC-MS/MS. For oocyte studies, 4 mice were hyperstimulated at 6 months post-chase and 7 mice at 10 months post-chase. A total of 104 and 45 oocytes were collected at 6 months and 10 months post-chase, respectively, and these samples were used for LC-MS/MS.

#### Ovary and oocyte collection

Ovaries were harvested and placed in a dish containing pre-warmed Leibovitz’s medium (L15) (Life Technologies Corporation, Grand Island, NY) supplemented with 3 mg/mL polyvinylpyrrolidone (PVP) (Sigma-Aldrich, St. Louis, MO), 0.5% penicillin-streptomycin (Life Technologies) (L15-PVP). Ovaries were either processed as described below for downstream analyses or used to isolate oocytes. Ovaries used to assess reproductive aging parameters were fixed for histological analysis in Modified Davidson’s (Electron Microscopy Sciences, Hatfield, PA) at room temperature for 2-4 hours with agitation and overnight at 4°C. Ovaries used for MIMS were cut in half and fixed in 2% glutaraldehyde (Electron Microscopy Sciences, Hatfield, PA) and embedded in LR white (EMS). Ovaries used for LC-MS/MS were snap frozen and stored at -80°C until use.

To maximize the yield of a synchronized population of fully grown oocytes for LC-MS/MS, mice were hyperstimulated with an intraperitoneal injection of 5 IU pregnant mare serum gonadotropin (PMSG) (Prospec Bio, East Brunswick, N). Ovaries were harvested 44-48 hours post-PMSG injection and placed in L15-PVP supplemented with 0.025% milrinone to maintain oocyte meiotic arrest (Sigma-Aldrich, St. Louis, MO).

Cumulus-oocyte-complexes (COCs) were released by puncturing antral follicles with insulin syringes. Oocytes were mechanically denuded from cumulus cells using a 75 μm stripper tip and washed thoroughly in L15/PVP/PS before being snap frozen. For analysis of ovulated eggs, mice were hyperstimulated as described above with PMSG followed by superovulation induction 44-46 hours later with an intraperitoneal injection of 5 IU human chorionic gonadotropin (hCG)(Sigma, St. Louis, MO). COCs were then collected from the oviducts of each mice 14-16 hours post-hCG injection, and the number of ovulated eggs were counted.

#### Ovarian follicle counts and histological analysis

The decline in the number of ovarian follicles is a hallmark of reproductive aging (*1*). Therefore, we evaluated follicle counts in histological sections of ovaries as done previously (*9*). In brief, fixed ovaries were washed three times with 70% ethanol and processed, dehydrated, and paraffin embedded using an automated tissue processor (Leica Biosystems, Buffalo Grove, IL). After embedding, ovaries were serial sectioned with every 5^th^ tissue section stained with hematoxylin and eosin (H&E). All H&E-stained tissue sections were digitally scanned at the University of Washington’s Histology and Imaging Core using Hamamatsu-HT imaging system (Hamamatsu Photonics, Hamamatsu City, Japan) at 20X magnification. Scanned images were uploaded and visualized using the NDP.view2 software (Hamamatsu Photonics, Hamamatsu City, Japan). Follicles were classified and counted in every 5^th^ tissue section. Follicles were classified by stage (primordial, primary, secondary, and antral) according to established criteria (*9*). Primordial follicles were comprised of an incomplete layer of squamous granulosa cells, while primary follicles contained a complete layer of cuboidal granulosa cells. Secondary follicles contained two or more layers of cuboidal granulosa cells. Antral follicles had more than eight layers of granulosa cells with the presence of an antrum, or fluid-filled cavity. All primordial and primary follicles were counted regardless of whether the oocyte’s nucleus was visible. Secondary and antral follicles were counted only if the nucleus was visible to avoid double counting. Only healthy follicles were included in the final counts. Atretic follicles containing abnormally shaped oocytes with dark, pyknotic granulosa cells were excluded. The average number of follicles per area of ovarian section was calculated and used to compare counts between each age cohort.

Ovarian fibrosis, characterized by excess collagen, is another hallmark of reproductive aging (*19*). We evaluated fibrosis in ovaries by Picrosirius Red (PSR), a histological stain which detects collagen I and III (*19*). Ovarian tissue sections were deparaffinized in Citrisolv (Fisher Scientific, Pittsburgh, PA) and rehydrated in 100%, 70%, and 30% ethanol baths. Slides were submerged in PSR staining solution composed of Sirius Red F3BA (Direct Red 80, C.I. 357.82, Sigma-Aldrich, St. Louis, MO) and picric acid (Sigma-Aldrich, St. Louis, MO) at 0.1% w/v for 40 minutes at room temperature. The slides were then incubated in acidified water made of 0.05 M hydrochloric acid (Fisher Scientific) for 90 seconds. Tissue sections were dehydrated in 100% ethanol baths, three times for 30-second incubations. After dehydration, slides were immersed in Citrisolv for five minutes and mounted with Cytoseal XYL (Fisher Scientific). PSR stained sections were then imaged with an EVOS FL Auto Imaging system (Thermo Fisher, Waltham, MA) using a 20X objective. Scans of whole ovarian tissue sections were performed to quantify the area of positive PSR staining using a threshold feature on ImageJ as previously described (Briley et. 2016). PSR positive staining was analyzed on two different ovarian tissue sections for three mice within each age cohort: 6 weeks, 6 months, and 10 months. Results were averaged to obtain average percent area of collagen.

#### Multi-isotope imaging mass spectrometry (MIMS) and data processing

Fixed LR white-embedded ovaries were sectioned to 0.5 microns and mounted on silicon wafers. At the Brigham and Women’s Hospital Center for NanoImaging, a NanoSims 50L (CAMECA Instruments Inc., Madison, WI) instrument was tuned to simultaneously measure ^12^C^14^N^-^, ^12^C^15^N^-^, and ^31^P^-^ secondary ions as described previously for imaging of a wide range of mouse and human tissues (*56*). Quantitative ^12^C^14^N images were used for histological representation of stereotypical ovarian structures and associated cell types. ^31^P images provided additional histological detail and were used for identification of nuclei due to the high phosphorus content of chromatin (*29*). Quantitative mass images for ^12^C^14^N and ^12^C^15^N were used to generate quantitative ^15^N:^14^N ratio images. For imaging of swathes of the tissue section, images were acquired in chain analysis mode of sequential adjacent fields (dimensions of 45 μm x 45 μm or 50 μm x 50 μm). Sequential tiles were stitched together to make mosaic images for visualizing large sections of the ovary (**Fig. 1B**). Some features were then imaged at higher resolution with smaller field sizes. All images were processed and further analyzed using the most recent version of the OpenMIMS 2.0 plugin (https://github.com/BWHCNI/OpenMIMS) to ImageJ (*57*). ^15^N-labeling was visualized by a hue saturation intensity (HSI) transformation of the ^12^C^15^N/ ^12^C^14^N ratio. The color scale of HSI images is set such that the lower blue bound of the scale is at the ^15^N natural abundance of 0.37% (expressed as 0 % above background). The upper magenta bound of the scale is set to reveal labeling differences. The quantitative isotope ratio measurements that form the basis for the images and that are used for statistical analyses are not affected by scaling changes that modify the visual appearance of the images (**fig. S2 A-B).** A combination of ^14^N and ^31^P images were used to manually select regions of interest within each ovarian section (e.g. individual cells or subcellular structures). Structures and cell types were identified based on morphology and their anatomic location. Cells that were not identifiable or were not visualized due to low ion counts were excluded from the analysis. Typical reasons for difficult identifying cells include sectioning artifacts (cracks, wrinkles) or certain cells that were located at the juncture between adjacent imaging fields, where there is often lower yield of secondary ions. This edge effect is seen to variable degrees in the mosaic images as dark regions at the periphery of an imaging field. All quantitative data for ^15^N-labeling was presented as the ^15^N/^14^N ratio (percentage above natural abundance).

#### Mass Spectrometry (MS) sample preparation: ovaries

Isolated ovaries were homogenized directly in 6M guanidine hydrochloride solution using bead-based Precellys 24 homogenizer, followed by processing with ProteaseMAX according to manufacturer’s protocol. Samples were reduced with 5 mM Tris(2-carboxyethyl)phosphine (TCEP; vortex 1 hr at RT) alkylated in the dark with 10 mM iodoacetamide (IAA; 20 min), diluted with 50 mM ABC and quenched with 25 mM TCEP. Samples were digested with sequencing grade modified trypsin overnight at 37°C with shaking, spun down (15,000 × g for 15 min at RT), placed in a new tube and acidified with TFA to a final concentration of 0.1%. A total of 100 μg of digested and acidified sample was fractionated using High pH Reversed-Phase Peptide Fractionation Kit (Pierce, Cat# 84868). Fractions were step eluted in 300 μL buffer of increasing acetonitrile (ACN) concentrations with decreasing concentration of Triethylamine (0.1%) as per manufacturer. Samples were dried down with vacuum centrifugation for future MS analysis.

#### MS sample preparation: GeLC/MS on oocytes

Isolated oocytes were lysed directly in RIPA buffer, mixed with 6X SDS sample buffer, boiled for 5 minutes, and separated by SDS-PAGE using 10% Tris-glycine gels (Thermo Scientific, Cat# XV00100PK20). Gels were stained using Oriole fluorescent gel stain solution, scanned using Bio-rad Chemidoc XRS system, and cut into sections, chopped into 1mm x 1 mm cubes, and processed for in-gel digestion. The gel separating oocytes collected at t=6 months was cut into 36 individual pieces, whereas the gel separating oocytes collected at t=10 months was cut into 24 pieces. Gel pieces were incubated in 10 mM TCEP (in 50 mM ABC; 1 hr at 37 °C). Liquid was replaced by 50 mM IAA (in 50 mM ABC; 45 min at RT in dark), followed by 50 mM TCEP (in 50 mM ABC; 30 min at RT). Gel pieces were washed with 50 mM ABC (3x) and digested with sequencing grade modified trypsin (1 μg in 50 mM ABC, overnight at 37 °C, with shaking). The following day, supernatant was collected into new tube and the gel piece were subjected to three rounds of incubations with 50% ACN and 5% FA solution (30 min at RT, with shaking). Supernatant was collected after each incubation, combined, and dried down with vacuum centrifugation. Samples were re-suspended in 0.5% TFA, desalted with Pierce C18 spin columns (Thermo Scientific, Cat# 89873) per manufacturer’s instructions, and dried down with vacuum centrifugation for future MS analysis.

#### MS analysis

Dried samples were re-suspended in 20 μL Buffer A (94.875% H2O with 5% ACN and 0.125% FA) and three micrograms, as determined by microBCA assay (Thermo Scientific, Cat# 23235) of each fraction or sample were loaded via auto-sampler with either Thermo EASY nLC 100 UPLC or UltiMate 3000 HPLC pump, onto a vented Pepmap 100, 75um x 2 cm, nanoViper trap column coupled to a nanoViper analytical column (Thermo Scientific) with stainless steel emitter tip assembled on the Nanospray Flex Ion Source with a spray voltage of 2000 V. A coupled Orbitrap Fusion was used to generate MS data. Buffer A contained 94.785% H_2_O with 5% ACN and 0.125% FA, and buffer B contained 99.875 ACN with 0.125% FA. MS parameters were as follows: Ion transfer tube temp = 300 °C, Easy-IC internal mass calibration, default charge state = 2 and cycle time = 3 s. Detector type set to Orbitrap, with 60K resolution, with wide quad isolation, mass range = normal, scan range = 300-1500 m/z, max injection time = 50 ms, AGC target = 200,000, microscans = 1, S-lens RF level = 60, without source fragmentation, and datatype = positive and centroid. MIPS was set as on, included charge states = 2-6 (reject unassigned). Dynamic exclusion enabled with n = 1 for 30 s and 45 s exclusion duration at 10 ppm for high and low. Precursor selection decision = most intense, top 20, isolation window = 1.6, scan range = auto normal, first mass = 110, collision energy 30%, CID, Detector type = ion trap, OT resolution = 30K, IT scan rate = rapid, max injection time = 75 ms, AGC target = 10,000, Q=0.25, inject ions for all available parallelizable time. For ovary samples the chromatographic run was 4.5 hours per fraction, with the following profile of Buffer B: 2% for 7 min, 2-7% for 1 min, 7-10% for 5 min, 10-25% for 160 min, 25-33% for 40 min, 33-50% for 7 min, 50-95% for 5 min, 95% for 15 min, then back to 2% for the remaining 30 min. For the GeLC/MS oocyte samples, the chromatographic run was 75 min per gel section with the following profile of Buffer B: 2-8% for 6 min, 8-24% for 10 min, 24-36% for 20 min, 36-55% for 10 min, 55-95% for 10 min, 95% for 10 min, then back to 2% for remaining 9 min.

#### MS Data Analysis and Quantification

Protein identification/quantification and analysis were performed with Integrated Proteomics Pipeline - IP2 (Integrated Proteomics Applications, Inc., San Diego, CA. http://www.integratedproteomics.com/) using ProLuCID (*58, 59*), DTASelect2 (*60, 61*), Census and QuantCompare. Spectral raw files were extracted into MS1, MS2 files using RawConverter 1.0.0.0 (http://fields.scripps.edu/downloads.php). The tandem mass spectra were searched against mouse database (downloaded on 03-25-2014). Searched spectra were matched to sequences using the ProLuCID/SEQUEST algorithm (ProLuCID version 3.1) with 50 ppm peptide mass tolerance for precursor ions and 600 ppm for fragment ions. ProLuCID searches included all fully and half-tryptic peptide candidates that fell within the mass tolerance window and had with unlimited mis-cleavages. Carbamidomethylation (+57.02146 Da) of cysteine was considered as a static modification. Peptide/spectrum matches (PSMs) were assessed in DTASelect2 using the cross-correlation score (XCorr), and normalized difference in cross-correlation scores (DeltaCN). Each protein identified was required to have a minimum of one peptide (-p1) of minimal length of six amino acid residues. False discovery rate (FDR) was set to 1% at the protein level, for all experiments. Peptide probabilities and FDR were calculated based on a target/decoy database containing the reversed sequences of all the proteins appended to the target database (*62*). Each dataset was searched twice, once against light (^14^N) and then against heavy (^15^N) protein databases, as described previously(*14*). In the light searches, all of the amino acid residues were considered to contain only ^14^N nitrogen, while in the heavy searches, all the amino acid residues were considered to contain only ^15^N nitrogen. After the results from ProLuCID were filtered using DTASelect2, and the assembled search result file was used to obtain quantitative ratios between ^14^N and ^15^N using the software Census (*30, 63*).

Long-lived proteins were identified as previously described, with modifications (*14*). Briefly, in order for a protein to be considered as long-lived, the protein had to be identified in our heavy/light search by at least one long-lived peptide (^15^N-peptide). Peptide ratio measurements were filtered in Census based on a correlation threshold, and only peptides with correlation coefficient above 0.5 were used for further analysis. For singleton analysis, we required the ^14^N/^15^N ratio to be greater than 5.0 and the threshold score to be greater than 0.5. Identified peptides were further filtered based on their average peptide enrichment (APE), which we set to 0.9, and peptide profile score, which was set to 0.8. Proteins were only identified as long-lived if they had more than three long-lived peptides that passed the above filtering (except for GeLC/MS experiments where one peptide was required). Fractional abundance were calculated according to the following formula: ^14^N values: FA = 100 – (100∗(1/(1+AR))), where FA = fractional abundance and AR = area ratio.

#### Gene Ontology Analysis

GO analysis was performed using the Pantherdb (*64*). The “query” is defined as proteins identified as long-lived in the analyzed tissue (based on ^14^N-peptide identification), and the reference is defined as all proteins identified in the same tissue analyzed (^14^N and ^15^N-peptide identification).

#### Statistical analysis

Statistical analyses were conducted using GraphPad Prism, version 9 (GraphPad Software, Inc.). A Student’s t-test was performed for comparisons between two groups. For comparisons of more than two groups, a one-way ANOVA was used. A one sample t-test was used to compare ^15^N/^14^N ratios of cytoplasmic and nuclear regions to a hypothetical value of 1. Values >1 signified a cell had greater ^15^N-labelling abundance in the nucleus compared to the cytoplasm. Values <1 represented cells with a lower ^15^N-labelling abundance in the nucleus compared to the cytoplasm. Values equal to 1 represented a cell with a ^15^N-labelling abundance that were equivalent in both nuclear and cytoplasmic regions. Data were considered significant with a p-value <.05 (* p-value <.05, ** p-value <.01, *** p-value <.001). Variability of groups was denoted as standard error of mean (SEM).

**Fig. S1:**
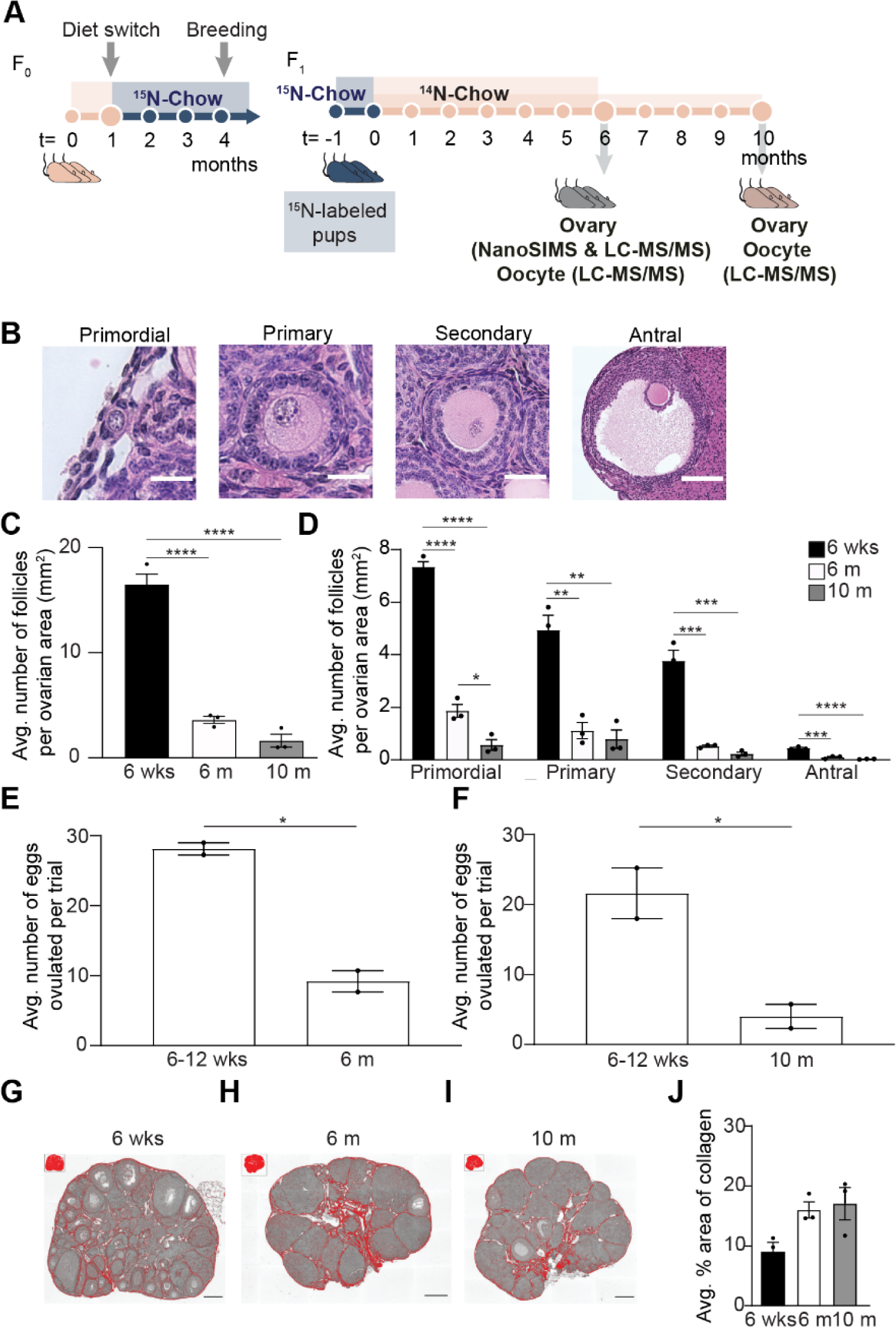
Multi-generational whole animal pulse-chase labeling design along the reproductive aging continuum. **(A)** Wild-type female FVB mice (n=3) were fed a ^15^N-diet for 13 weeks and were maintained on a ^15^N-labeled diet through breeding, pregnancy, and weaning to produce ^15^N-labeled pups. Labeled pups were sacrificed after switching over to a ^14^N-diet for a chase period of 6- or 10-months. Ovaries and oocytes were designated for NanoSIMS, or liquid-chromatography/mass spectrometry. Female FVB mice experience age-associated changes in ovarian reserve, ovarian microenvironment, and gamete quality. **(B)** Representative images of each follicle class from mice of 6 weeks old and 6 months old. **(C)** Average follicle number per area of ovarian section from mice of the following ages: 6 weeks, 6 months, and 10 months (N= 3 mice per age cohort). **(D)** Graph represents average number of follicles within each follicle class per area of ovarian section for mice ages 6 weeks, 6 months, and 10 months. (**E**) Comparison of average number of eggs ovulated per trial for mice at 6-12 wks, 6 months, and (**F**) 10 months. The data points represent the average number of eggs collected per mouse from two independent trials. In each trial, oviducts from 3-4 mice were pooled per age group. Representative processed color threshold images of PSR-stained ovarian tissue sections from mice **(G)** 6 weeks old, **(H)** 6 months old, and **(I)** 10 months old**. (J)** Graph comparing the average percent area of PSR-positive staining per ovarian section (pixels/μm^2^). Data are shown as mean ± SEM. Statistical analysis was performed using a one-way ANOVA. Asterisk denotes statistical significance (* p≤.05; ** p≤.01; *** p≤.001; **** p≤.0001). Scale bar for images of (B) primordial, primary, secondary, and antral follicles are 20 μm, 40 μm, and 140 μm, respectively. Scale bars in (G-I) are 180 μm.

**Fig. S2:**
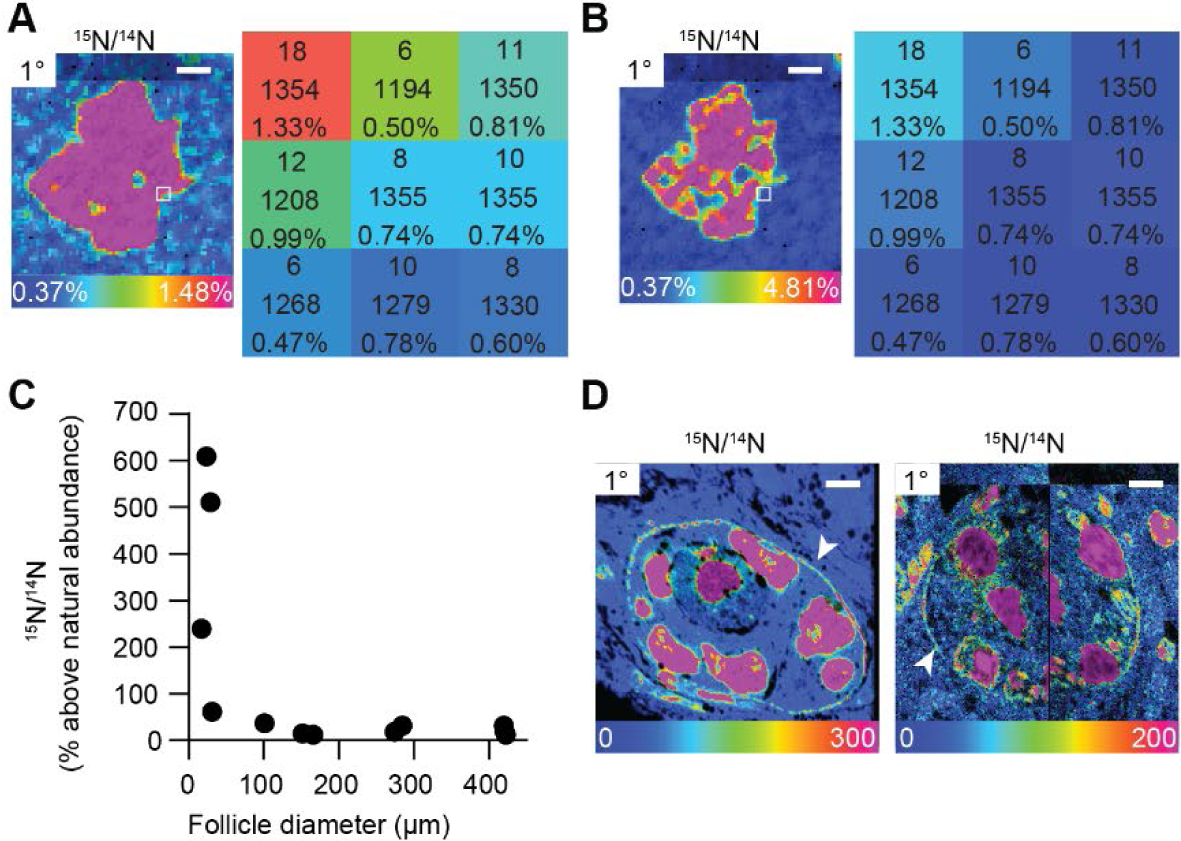
Multi-isotope imaging mass spectrometry (MIMs) uncovers structures enriched with ^15^N. (A) Hue saturation intensity image maps ^15^N/^14^N ratio across nuclear region of a primary follicle. Using a rainbow scale, blue is set to the natural ratio of ^15^N (0.37%) and overabundance is set to 1.48% (or 300% above the natural ratio). Each pixel provides quantitative information. The numbers in each pixel represent the number of ^15^N ions, ^14^N ions, and the ^15^N/^14^N ratio, respectively. (B) Changes to the rainbow scale can be used to emphasize regional ratio differences and change the visual representation of the data. Visual changes to HSI images due to different scales, does not change the quantitative data behind each pixel. (C) Total ^15^N abundance of each follicle plotted by follicle diameter shows smaller follicles containing high abundance of ^15^N. (D) HSI images reveal high ^15^N abundance concentrated at the basement membrane of early-stage follicles. Scale bar (panel A-B) =2.5 μm, (D) left= 4.0 μm, (D) right = 4.5 μm.

**Fig. S3:**
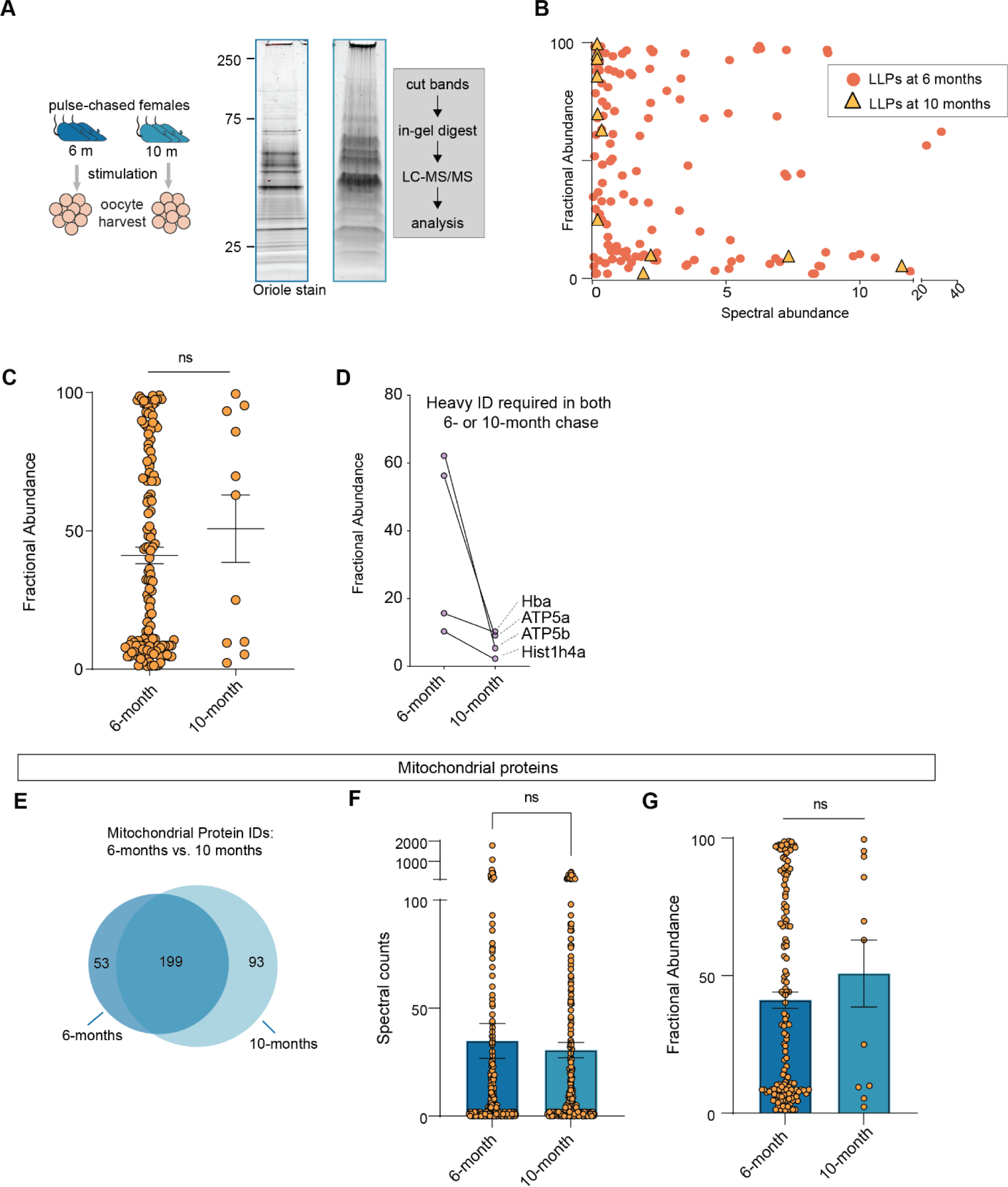
Long-lived proteins at 6- and 10-months chase points in oocytes. **(A)** Experimental scheme to identify and measure LLPs mouse oocyte. **(B-C)** Fractional abundance of each identified LLPs at both 6 months and 10 months as compared to the spectral abundance for each LLPs. **(D)** Plots showing decrease in FA values for same proteins identified as LLPs at 6 and 10-month chase. (E) Venn diagram illustrating the overlap of mitochondrial proteins identified at 6 and 10-month time points (F) Spectral counts of mitochondrial proteins as well as the (G) fractional abundance of each identified mito-LLPs at both 6 months and 10 months, illustrating that even though there are no significant difference in spectral counts at each timepoint, the number of quantified proteins decreased dramatically between 6 and 10 months of chase.

Table S1. Summary of proteins identified in 6- and 10-month chase ovaries.

Table S2. Summary of proteins identified in 6 and 10-month chase oocytes.

